# PEDOT:PSS-Modified Cotton Conductive Thread for Mass Manufacturing of Textile-Based Electrical Wearable Sensors by Computerized Embroidery

**DOI:** 10.1101/2021.12.06.471240

**Authors:** Fahad Alshabouna, Hong Seok Lee, Giandrin Barandun, Ellasia Tan, Yasin Çotur, Tarek Asfour, Laura Gonzalez-Macia, Philip Coatsworth, Estefanía Núnez-Bajo, Ji-Seon Kim, Firat Güder

## Abstract

The textile industry has advanced processes that allow computerized manufacturing of garments at large volumes with precise visual patterns. The industry, however, is not able to mass fabricate clothes with seamlessly integrated wearable sensors, using its precise methods of fabrication (such as computerized embroidery). This is due to the lack of conductive threads compatible with standard manufacturing methods used in industry. In this work, we report a low-cost poly(3,4-ethylenedioxythiophene) polystyrene sulfonate (PEDOT:PSS)-modified cotton conductive thread (PECOTEX) that is compatible with computerized embroidery. The PECOTEX was produced using a crosslinking reaction between PEDOT:PSS and cotton thread using divinyl sulfone as the crosslinker. We extensively characterized and optimized our formulations to create a mechanically robust conductive thread that can be produced in large quantities in a roll-to-roll fashion. Using PECOTEX and a domestic computerized embroidery machine, we produced a series of wearable electrical sensors including a facemask for monitoring breathing, a t-shirt for monitoring heart activity and textile-based gas sensors for monitoring ammonia as technology demonstrators. PECOTEX has the potential to enable mass manufacturing of new classes of low-cost wearable sensors integrated into everyday clothes.

## 1. Introduction

Wearable sensors offer the possibility of monitoring health continuously using noninvasive devices attached to the body. This approach allows long-term monitoring of biophysical and (bio)chemical signals related to health and well-being, ranging from monitoring of activity, heart rate and breathing to levels of glucose, lactate or ions in sweat.^[1–4]^ Continuous measurements provide an entirely new way to study the health of an individual by comparing them of today to themselves in the past.^[5]^ This is particularly important for diagnosing diseases early and monitoring the effectiveness of treatments to improve outcomes. One of the main bottlenecks for emerging wearable sensors is that they often require the user to wear and carry around additional hardware that is needed for the measurements which can come in the form of watches, tattoos, patches and straps among others.^[6–10]^ There is, of course, a practical limit to how many dedicated and standalone instruments one can wear on any given day; this limitation slows the adoption of many wearable innovations.

Textile fabrics are worn by everyone throughout the world as everyday clothes. Although clothes are in constant contact with the body, sensors are rarely incorporated into clothing in real-world applications. There are at least three big unaddressed challenges concerning the seamless integration of electrical and electrochemical sensors in clothing: materials and components used to produce sensors *(i)* alter the physical and chemical properties (*e.g.,* breathability, wettability, feel-on-the-skin) of fabrics; *(ii)* are generally not compatible with existing, well-established methods of fabrication used by the textile industry which predominantly rely on mechanically robust and durable threads/fibers and *(iii)* are vulnerable to cracking, delamination and chemical degradation during use or cleaning.^[11, 12]^

The textile industry has substantially computerized their manufacturing processes to increase the volume of production while reducing time and cost. To accelerate the adoption of new electrical sensing technologies by the textile industry, electrical sensing elements and interconnects must be compatible with the existing methods of manufacturing and should not require additional infrastructure or new high-cost instruments. Because mechanically robust organic fibers, yarns and threads, such as cotton, polyester and nylon, are the standard materials in the textile industry, there have been attempts to produce conducting or semiconducting versions of these materials through the application of organic and metallic coatings.^[13, 14]^ Yarns produced by coatings require complex synthesis and may use toxic compounds in the process;^[15–17]^ most importantly, however, surface coatings applied are generally fragile and cannot withstand the mechanically harsh abrasive processes used in the manufacturing of textiles. Because of this, conductive organic yarns reported to date have either been exclusively hand embroidered to incorporate sensors into fabrics or required highly specialized and expensive instruments.^[18–20]^

Computerized embroidery would allow the transfer of sensing structures with high spatial definition, designed on a computer, to fabrics in an accelerated fashion. The lack of conductive and semiconductive yarns compatible with computerized embroidery has particularly been a big issue to create sensors on fabrics reliably. Among all conducting polymers, the aqueous suspension of PEDOT:PSS has been widely used in textile coating applications due to its processability and biocompatibility.^[21]^ Müller and coworkers have reported computerized embroidery of all organic conductive silk yarns coated with PEDOT:PSS.^[22]^ Their material exhibited high electrical conductivity (70 S cm^-1^) before embroidery but upon embroidery, the conductivity of the yarns likely dropped significantly due to abrasion (note that the authors did not specify exactly how much this drop was).^[22]^ In another work by the same group, yarns of regenerated cellulose derived from wood pulp were coated with PEDOT:PSS, for some samples with Ag nanowires, and used in machine sewing of thermocouples for energy harvesting.^[23]^ The authors used their yarn only in the bottom thread (or bobbin thread) configuration which sacrifices spatial definition for reduced tension and mechanical requirements for the thread. The bottom thread configuration is not commonly used for embroidering patterns by the textile industry due to poor appearance. Furthermore, the authors still relied on conductive silver paste added manually to connect thermocouples together to minimize resistive losses. Although Müller and coworkers have made progress toward all organic conductive threads, mechanical robustness and high costs of threads remain to be unsolved problems for fabricating sensors through computerized embroidery.

In this work, we report electrically conductive and mechanically robust all organic cotton-based low-cost threads that are fully compatible with domestic and industrial computerized embroidery machines. We solved the problem of mechanical robustness of conductive threads during embroidery by exploiting a crosslinking reaction using divinyl sulfone (DVS) as a crosslinker in a roll-to-roll reaction involving poly(3,4-ethylenedioxythiophene) polystyrene sulfonate (PEDOT:PSS), ethylene glycol (EG) and the cellulose substrate (**Figure 1**). PEDOT:PSS-modified conductive cotton threads (PECOTEX) produced were structurally, mechanically, electrically/electrochemically characterized and used for the seamless integration of wearable electrical sensors into clothing. We produced facemasks for monitoring breathing, a t-shirt for monitoring heart activity and textile-based gas sensors for monitoring ammonia as technology demonstrators.

**Figure 1.**
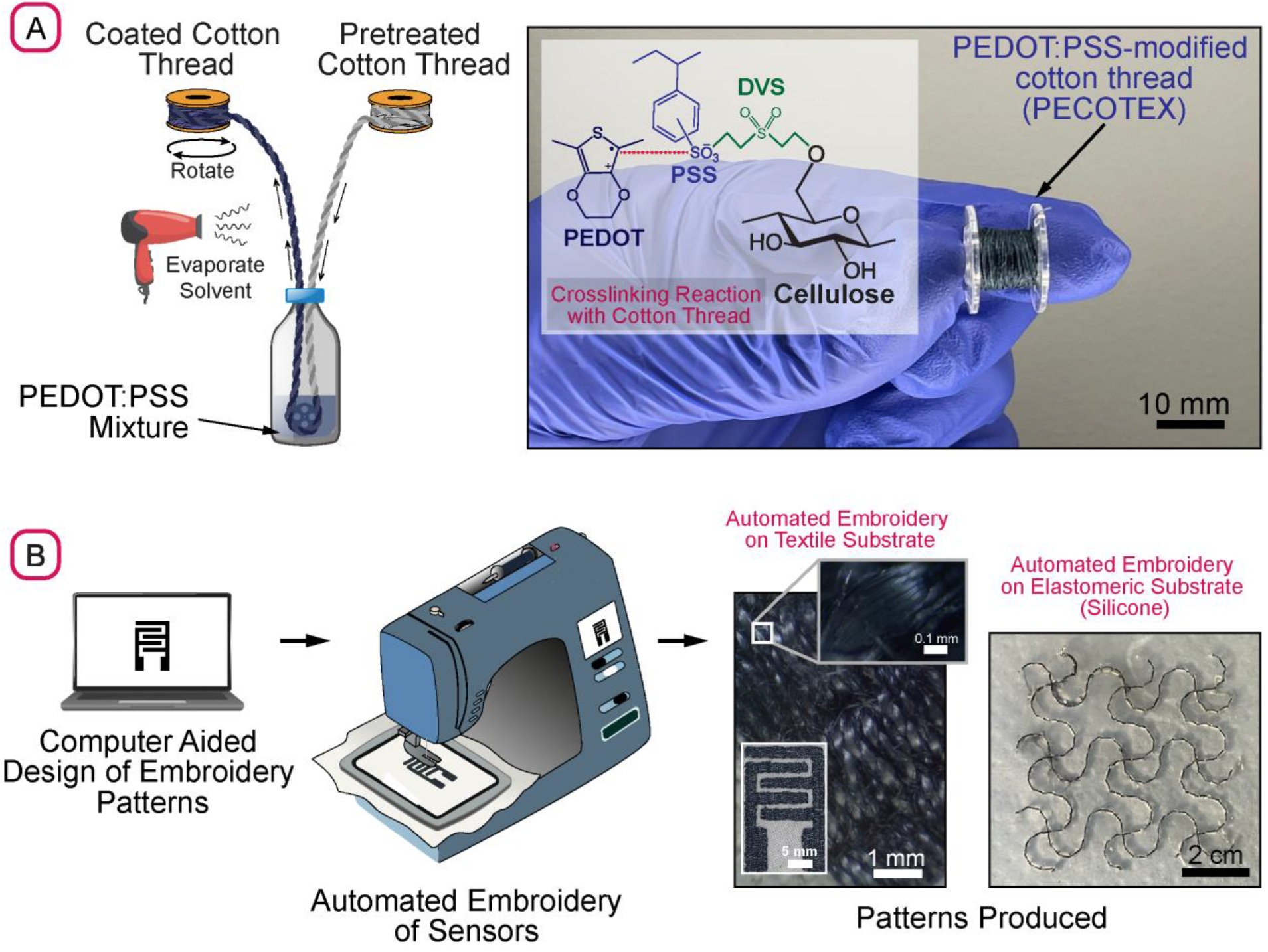
**(A)** Schematic illustration of synthesis of the PEDOT:PSS-modified cotton thread (PECOTEX) and a photograph of the produced thread on bobbin. The inset shows the chemical structure of the product of the crosslinking reaction between PEDOT:PSS and cellulose surfaces within the cotton thread; **(B)** Process flow for the computerized embroidery of patterns using PECOTEX on cotton textile (interdigitated electrodes) and silicone (serpentine structures) substrates.

## 2. Results and Discussion

### 2.1. Synthesis of PEDOT:PSS-based Conductive Threads

Before dying the cotton thread with the PEDOT:PSS-based conductive ink formulation, we pretreated the threads in an aqueous solution of a scouring detergent, METAPEX 38 (manufactured by ArtVanGo, UK) stirred at 80 °C for 2 h. This was followed by oxidative desizing for 5 h using bleach (Waitrose, UK) to remove the cuticle layer which consists mostly of fats and waxes (see Supporting Information **Figure S1** for more information on pretreatment). The threads were then rinsed several times in deionized (DI) water to clean the excess reagents and impurities. Pretreatment of the cotton thread is important to improve the adhesion of the conductive dye to the surface by increasing the number of reactive sites on the fibers which helps to produce a uniform and denser coating. After pretreatment, the cotton threads were dyed using a homemade, continuous roll-to-roll coating system (Figure S1) which was digitally controlled by an Arduino microcontroller connected to a servomotor (Figure 1A). This setup allowed passing a bobbin of thread through a sequence of vials containing liquid reagents/ink with adjustable speed. A standard hairdryer (Red Hot Professional Hair Dryer) was used to blow hot air from a distance (~15 cm) to remove any excess water from the thread before winding on the bobbin to prevent sticking. With this setup, we were able to produce 100 m of thread over a period of six hours in our laboratory in an automated fashion. Using the roll-to-roll coater, we produced six different types of threads (**Figure 2A**) with different formulations, all of which included an aqueous dispersion of PEDOT:PSS as the conductive dye with reagents added to improve the electrical and physical characteristics of the final coating: *(i)* Type I thread was only coated with PEDOT:PSS. *(ii)* Type II included PEDOT:PSS + 5% wt. Dimethyl Sulfoxide (DMSO). (*iii*) Type III thread consisted of PEDOT:PSS + 5-10% wt. EG. *(iv)* Type IV contained PEDOT:PSS + 3-10% DVS. *(v)* Type V included PEDOT:PSS + 5% wt. DMSO + 3% DVS. *(vi)* Type VI consisted of PEDOT:PSS + 3-7% wt. EG + 3-7% DVS. We chose DMSO and EG, widely studied cosolvents, as additives to improve the conductivity of PEDOT:PSS.^[24]^ Since our eventual goal is to create mechanically robust conductive threads, that are compatible with computerized machine embroidery, to further improve the adhesion of PEDOT:PSS to the OH-rich surfaces of the cellulose fibers, we also experimented with the crosslinker DVS. DVS has two vinyl reactive ends that can replace the hydrogen in OH groups, present on the cellulose surface and PSS by vinyl sulfone fragments via Oxa-Michael nucleophilic addition. DVS could, therefore, covalently crosslink different elements within the coating while also forming covalent bonds with the cellulose surface, similar to vinyl sulfone reactive dyes to textiles ^[25]^, to yield a robust conductive coating (Figure 1A - inset). Furthermore, unreacted DVS acts as a second dopant to improve the conductivity of PEDOT, further increasing its utility as an additive.^[26]^

**Figure 2.**
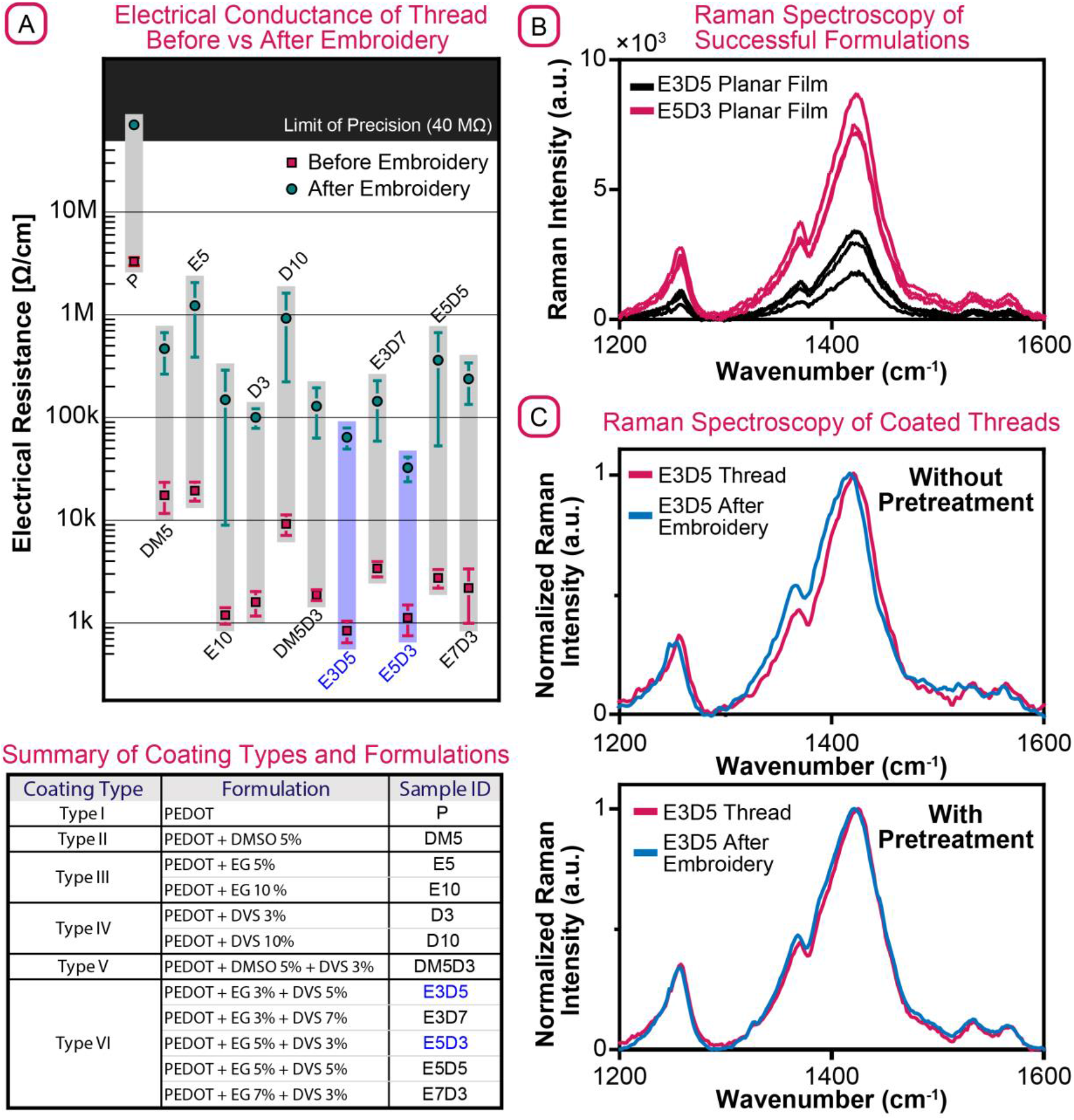
**(A)** Electrical resistance of the conductive threads with different formulations before and after embroidery. Table provides a summary of all formulations. Error bars: SD, n=5. **(B)** Raman spectra for planar films of the two successful formulations (E5D3, E3D5). E3D5 was eventually selected because of smaller amount of dedoped PEDOT present in the film. **(C)**. Normalized Raman spectra of E3D5 with and without pretreatment. Pretreatment increases -OH reaction sites on cellulose, hence the robustness of the coating.

### 2.2. Optimization of Formulation of Coating

To find the best formulation for a robust, PEDOT:PSS-based electrically conductive coating for cotton threads compatible with computerized embroidery and understand the role of each additive, we have performed a series of experiments. Because the most important characteristic for our threads is the electrical resistance (1/ electrical conductance), we measured the electrical resistance of each thread created with a different formulation before and after embroidery (Figure 2A). The ideal thread for computerized embroidery should have the lowest electrical resistance before and after embroidery; to test the threads, a bobbin of the conductive thread was placed into the embroidery machine as top bobbin (**Figure S2**) and square line stitches (stitch length 2 mm) of 25 cm^2^ were embroidered onto a cotton textile using a domestic computerized embroidery system (Diamond Royale™ by HUSQVARNA VIKING ®). We chose a square pattern because the abrasive forces exerted on the thread primarily originate in the eye of the needle and the magnitude of abrasion forces depends on the direction of embroidery. After embroidery, the embroidered patterns were unstitched, electrical resistance was measured using a multimeter and compared to before embroidery for each formulation (Figure 2A). For all samples regardless of the formulation, the electrical resistance of the threads increased after embroidery due to the mechanical abrasion during stitching which breaks the conductive paths and partially removes the coating (**Video S1**); however, while some formulations, such as Type I thread (*i.e.*, coated with PEDOT:PSS only), exhibited a large starting resistance and a large increase in resistance after embroidery (electrical resistance before embroidery: 3.3 MΩ cm^-1^, electrical resistance after embroidery: > 40 MΩ cm^-1^ – *i.e.*, beyond the limit of precision of the multimeter), Type VI thread – formulations consisting of PEDOT:PSS + 5% EG + 3% DVS (sample ID: E5D3) and PEDOT:PSS + 3% EG + 5% DVS (sample ID: E3D5) – had an electrical resistance of 800 −1000 Ω cm^-1^ before embroidery and 10 – 80 kΩ cm^-1^ after embroidery. In these experiments, we have also noted that the addition of EG and DMSO also improved conductance of the threads before and after embroidery because of the uncoiling and stretching effect of the PEDOT chains which brings PEDOT molecules closer together facilitating the charge transport (Figure S2). Different to DMSO, EG, however, is also known to increase the adhesion of PEDOT:PSS to hydroxyl (OH)-rich surfaces by hydrogen bonding.^[22]^ Increasing the concentration of the EG enhanced the conductivity of thread before and after embroidery as the hygroscopic characteristics of EG also allows more water content to remain on the thread. The crosslinker DVS, when used on its own, performed better than PEDOT:PSS alone, but made the threads stiffer and more fragile especially at higher concentrations. Hence, a combination of PEDOT:PSS, EG and DVS performed the best due to enhanced hydrogen bonding and crosslinking between PEDOT:PSS, EG and the surface. The reaction of DVS with EG also produces a rubbery polymer as reported previously by Mantoine and coworkers^[26]^ which improves the flexibility of the thread during embroidery hence produces a more mechanically durable coating.

In order to select the most optimum formulation between E5D3 and E3D5, we performed Raman spectroscopy for more detailed chemical and structural analysis of PEDOT, the conductive component in the coating (**Figure 2B** and **2C**). To eliminate the influence of the microstructure of the cellulose thread on the intensity of the signals, we first produced freestanding films in Petri dishes and performed the measurements (Figure 2B). The experiments showed that E5D3 produced a stronger spectral intensity than E3D5 across the board. This result indicates that a larger amount of PEDOT in E5D3 is in resonance with the excitation wavelength hence, in a more dedoped (neutral) state than E3D5. This means that E5D3 is less conductive than E3D5, yet the former showed lower measured electrical resistance than the later as in Figure 2A. The difference in electrical resistance was associated with EG content, EG is hygroscopic by nature which means E5D3 contained temporarly more water molecules, hence was more conductive during the measurement. The addition of EG plays a significant role in increasing the conductivity through the conformational change of PEDOT chains from coiled to liner, but the excess of EG (>3wt % ^[26]^) increases the distance between the isolated PEDOT molecules leading to reduced charge transport (Figure S1). We, therefore, selected E3D5 for the rest of the experiments as the conductive coating for the cellulose thread.

We compared the normalized Raman intensities of E3D5 films coated on the cotton threads before and after embroidery and with and without pretreatment of the cellulose surface (Figure 2C). The main features in this comparison are the relative intensity and broadening of the full width half maximum (FWHM)^[27]^. The spectra were normalized to C=C (located at 1425 cm^-1^), so the relative intensity is the change in intensity of C-C intra peak (located at 1368 cm^-1^). Without pretreatment of the cotton thread, the embroidery process had a significant effect on the PEDOT structure with a 24% increase in the relative intensity. This indicates a conformational change in the PEDOT structure as π electron density is shifted towards the intra-ring C-C bonds.^[28]^ Additionally, there was a 4% broadening of the FWHM of the C=C peak. These changes indicate that there is a higher distribution of PEDOT conformations, hence a greater conformational disorder of the PEDOT segments was caused by the embroidery process. This suggests that PEDOT molecules were likely not anchored well on the surface of the fibers during coating which led to PEDOT structural changes during embroidery. In contrast, when the conductive coating was deposited on cotton threads with oxidative pretreatment, there was a lower degree of conformational change to the PEDOT segments after embroidery. The change in intra-ring C-C relative intensity was reduced to 7%, while the C=C peak FWHM was 1%. This observation agrees with the notion that pretreatment increases the number of -OH reaction sites on cellulose, therefore, improves the robustness of the coating during embroidery. Finally, we studied the impact of aging of E3D5 before and after embroidery by comparing normalized Raman spectra obtained after 24h and seven days which remained unchanged over this period, indicating short-term stability (**Figure S3**).

To further enhance the consistency of the coating, we coated the thread with four consecutive cycles of E3D5 formulation to produce PECOTEX. The electrical resistance of PECOTEX before embroidery improved to ~400 Ω cm^-1^ after four cycles of coating. The electrical resistance after embroidery, however, did not change as most of the excess PEDOT:PSS was abrasively removed during embroidery.

### 2.3. Electro-Mechanical and Microstructural Characterization of Threads

In order to quantify the mechanical robustness of PECOTEX, we measured the tensile strength of threads that have been pretreated and coated with the conductive coating and compared the results to cotton threads (no coating) with and without pretreatment (**Figure 3A** and **S4**). All threads tested could stretch approx. 20 mm and could withstand loads around 10 N. We noted, however, that the oxidative pretreatment process reduced the fracture strength of the threads up to 30%, likely due to the oxidative damage to the thread during pretreatment. The degree of damage could potentially be reduced by optimizing the pretreatment process. In any case, during embroidery, we did not observe snapping of the threads in our domestic computerized embroidery machine, suggesting that our coating process did not weaken the threads more than the loads experienced in a domestic embroidery machine to cause rapture.

**Figure 3.**
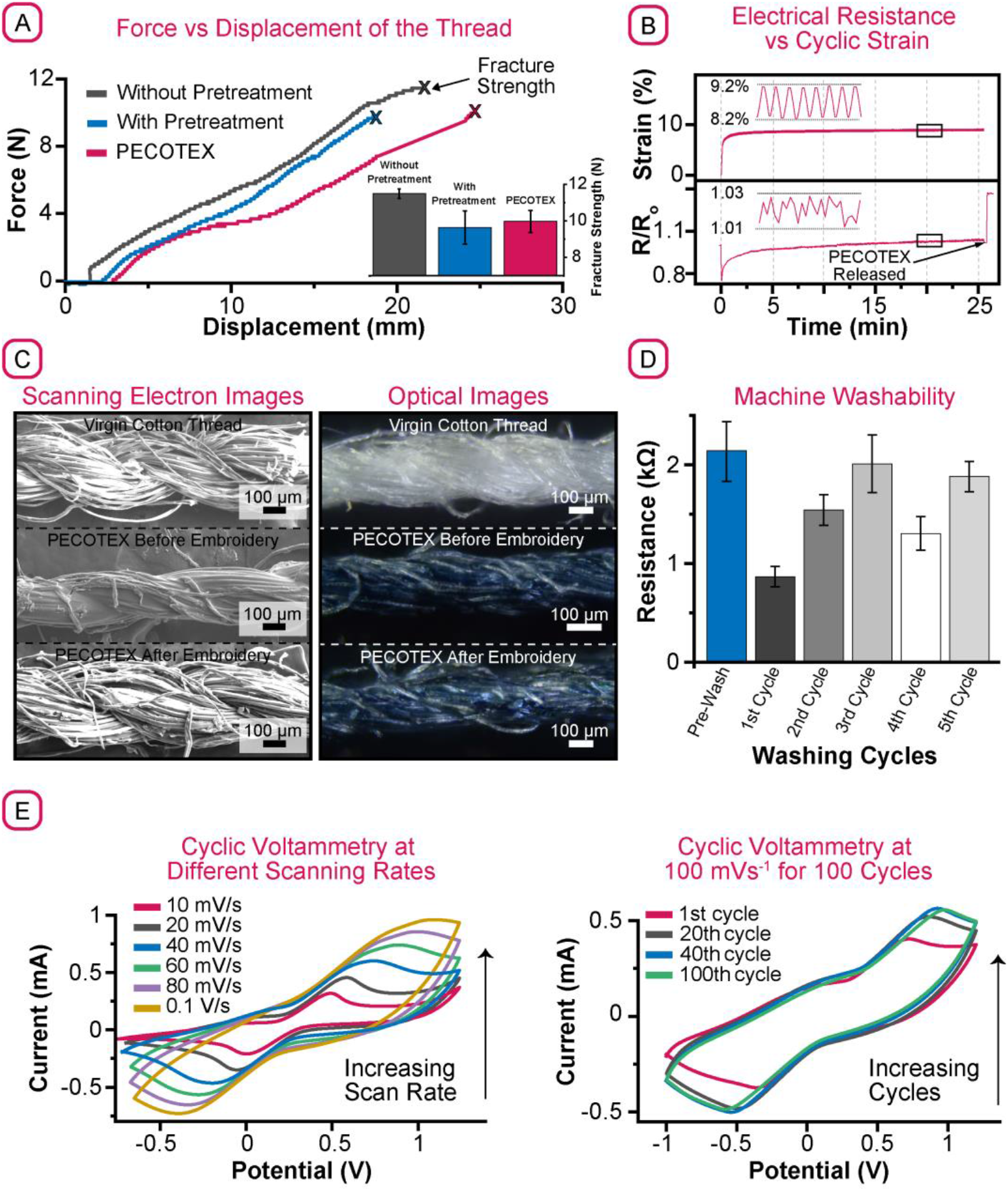
**(A)** Force-Displacement characteristics of the threads. Pretreatment weakens the thread, reducing its fracture strength; **(B)** Change in resistance of PECOTEX under cyclic strain up to 10%; **(C)** Scanning electron and optical micrographs of a cotton thread, PECOTEX and PECOTEX after embroidery; **(D)** Machine washability test for embroidered PECOTEX patterns up to five cycles. The resistance remained largely unchanged over the experiment. **(E)** Cyclic voltammograms of PECOTEX with increasing scan rate and number of cycles of scans. The PECOTEX is stable and suitable for most electrochemical sensing applications.

We characterized the electrical properties of PECOTEX under cyclic tensile loading (**Figure 3B** and S4). In this experiment, the threads were placed in a tensile tester and applied a periodically varying load between 2.5 N and 1.5 N with a frequency of 0.3 Hz for 1000 cycles. During this test the electrical resistance of the thread initially dropped likely due to the compression of the fibers together, increasing the number of parallel conductive paths within the thread. With repeated application of loading, the resistance increased slowly as the conductive pathways were damaged due to cyclic strain. When the thread was released, its resistance increased ~40%, indicating the fibers that were compressed together were now released, disrupting the parallel conductive pathways and the damaged coating increased the overall resistance. We did not observe a cyclic change in the resistance of the threads with the application of cyclic loading, however. These results suggest that the conductive coating was robust enough to withstand cyclic loading without a large impact on its resistance. These results also suggest that these fibers are not suitable for reversible strain sensing applications but could be utilized as single use strain sensors.

To study the impact of embroidery and harsh mechanical conditions experienced by the threads, we have visually inspected the microstructure of PECOTEX before and after embroidery using scanning electron and optical microscopy (**Figure 3C**). After the coating process, the diameter of PECOTEX decreased markedly in comparison to virgin cotton threads as the fibers were bound together by the coating - the surface of the coated thread appeared smoother. The polymer coating also rendered the threads blue due to the intrinsic color of PEDOT. The diameter of the coated threads increased almost back to the same diameter as the original virgin cotton thread because of the mechanical forces applied during embroidery which removed some of the coating and unbound the fibers. The partial removal of the coating and structural damage after embroidery were also evident in the optical images of the thread which appeared to have a slightly lighter tone of blue; these observations agreed with the results of electrical characterization of the threads before and after embroidery (Figure 2A) showing an increase in resistance after embroidery. Although there was a clear microstructural contrast between the PECOTEX before and after embroidery, the microstructure of PECOTEX after embroidery and virgin cotton thread was similar; PECOTEX could also be embroidered just as easily as the virgin thread – *i.e.,* the polymer coating did not render the cotton thread more sticky and less machinable.

Because most clothes are machine washed and reusable, we also studied the machine washability of PECOTEX after embroidery (**Figure 3D**) following the protocol described by the American Association of Textile Chemists & Colorists (AATTS).^[29]^ We machine embroidered 20 × 2 mm^2^ rectangular stitch patterns on a cotton textile substrate using the top-bobbin configuration and placed the patterns produced (n=6) in a washing bag (**Figure S5**). We measured the electrical resistance of the embroidered patterns after five cycles of washing in a domestic washing machine (Candy GOW475 Washer Dryer). We dried the samples in an oven at 50 °C for 1h between each wash cycle to remove all water from the samples before their electrical resistance was measured using a multimeter. After the first cycle of washing, the resistance of the embroidered patterns dropped ~50%, most probably due to the removal of some of the non-conducting molecule, PSS from the coating. After the 2^nd^ cycle of washing, the electrical resistance of the embroidered samples increased likely due to the mechanical damage caused by machine washing to the coating; however, after the 2^nd^ cycle of washing the electrical resistance of the patterns remained relatively stable (never exceeding their original resistance) for the rest of the cycles, suggesting that they are compatible with machine washing.

We have also characterized the electrochemical properties of the PECOTEX since one of the application areas targeted with this technology is wearable electrochemical sensing. For that, a three-electrode electrochemical cell was constructed consisting of a 3 cm long PECOTEX as the working electrode (WE) which was attached to a commercial screen-printed electrode strip (DropSens, Metrohm) with a Ag/AgCl reference (RE) and C counter electrode (CE). In a solution of 20 mM ferrocyanide/0.1 m KCl, we first performed cyclic voltammetry (CV) at different scan rates (10, 20, 40, 60, 80, 100 mV s^-1^) using ferrocyanide as an electrochemical probe (**Figure 3** and **S6**). We observed that the relation between the peak current (ip) and the square root of the scan rate (*v*^1/2^) was linear (**Figure S7**) as expected for freely diffusing redox species according to Randles-Sevcik equation. This linear relationship indicates that the rate of electron transfer of species in solution is diffusion-limited and not limited by the material characteristics of the electrode.^[30]^ This feature is important for the electrodes in electrochemical sensing in addition to high electrical conductance and chemical inertness. The peak-to-peak potential separation was observed to be higher than the ideal 59.2 mV, indicating a quasi-reversible oxidation/redox process.^[31]^

The conducting polymer was sufficiently stable for 100 cycles (**Figure 3C**) when measured at 100 mV s^-1^, showing an increase in the peak currents after 20 cycles. This was probably due to the exposure of the probe to the evolving microstructure of the electrodes stabilized after continued cycling; PEDOT:PSS-based electrodes are known to change their microstructure when polarized in solution.^[32]^

### 2.3. Fabrication of Wearable Sensors by Computerized Embroidery

To demonstrate the application of PECOTEX produced in this work, in wearable sensing, we have fabricated three different classes of proof-of-concept devices (**Figure 4)** including: *(i)* a garment (in this case a t-shirt) with embroidered electrodes for measuring the electrical activity of the heart by electrocardiography (ECG); *(ii)* a disposable mask containing embroidered electrodes for monitoring respiration electrically and; *(iii)* embroidered gas sensors for sensing NH_3_.

**Figure 4.**
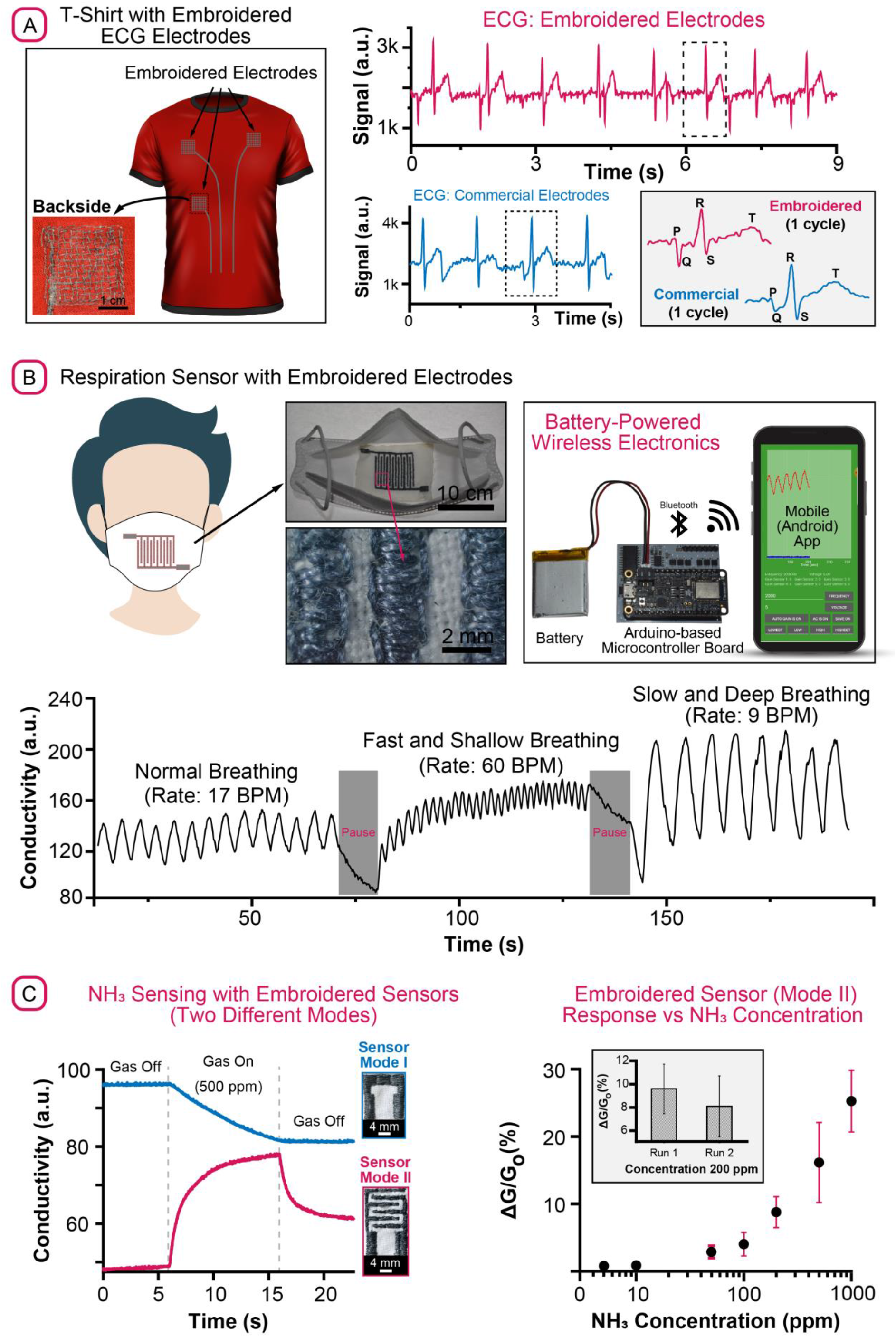
**(A)** Schematic of a t-shirt with embroidered PECOTEX ECG electrodes for monitoring cardiac activity. PECOTEX and commercial ECG electrodes produce clear and comparable waveforms that contain all medically relevant information. The inset shows the side of the electrode in contact with the skin; **(B)** Schematic of a disposable facemask containing embroidered PECOTEX electrodes for respiration sensing. The breathing patterns were captured using custom-made, battery-powered portable electronics and transmitted to a nearby mobile device for monitoring. Different respiratory activity was captured using the PECOTEX-based respiration sensor with low noise. **(C)** The electrical conductivity (normalized to 100%) of the PECOTEX-based embroidered gas sensors with two different sensing modalities when exposed to 500 ppm of ammonia at 70% relative humidity. The average change in electrical conductivity of the Mode II Sensor over a range of concentrations of NH_3_ (5 – 1000 ppm) for n=3 which shows an increasing response with concentration. The inset shows the response of PECOTEX-based embroidered Mode II Sensors (n=3) when exposed to 200 ppm of NH_3_ in two separate runs for the same sensors.

#### Continuous cardiac monitoring with an ECG t-shirt

Using computerized embroidery, we embroidered three electrodes (2.5×2.5 cm) on a textile t-shirt using PECOTEX for continuous monitoring of cardiac activity by ECG (**Figure 4A**). ECG is an important tool routinely used in medicine for diagnosing and monitoring cardiovascular conditions but often requires continuous monitoring for extended periods of time, ranging from weeks to months.^[33]^ We tested the ECG t-shirt for monitoring the electrical cardiac activity of a volunteer from our research group. After the volunteer put on the t-shirt, the embroidered electrodes were wetted with a conductive gel to improve the electrical contact with the skin - note that the gel penetrates the cotton t-shirt and electrodes providing excellent electrical contact (**Video S2**). We connected the embroidered electrodes to a microcontroller-based wireless ECG monitor and captured the cardiac activity of the volunteer while resting (**Figure S8**). After digital signal processing (**Figure S9** and **S10**), the ECG waveforms captured by the embroidered electrodes were comparable to the waveforms captured by the commercial electrodes (Figure 4A) in terms of noise artefacts. In both waveforms, the characteristic weaves such as the Q, R, S etc. were clearly identifiable allowing detecting of the heart rate and other important medical metrics, demonstrating that the ECG t-shirt could potentially be used for monitoring cardiac activity continuously.

#### Electrical monitoring of respiration with an embroidered sensor

We have produced embroidered electrical respiration sensors on cotton textile substrates and embedded them in disposable facemasks (used commonly by the public during the COVID-19 pandemic) for continuous monitoring of respiratory activity (**Figure 4B**). The sensor consisted of two embroidered electrodes used for measuring the ionic conductivity of the textile substrate; the ionic conductivity of the highly hygroscopic cotton textile changed during the cycles of inhalation and exhalation which modified the level of moisture adsorbed within the substrate.^[34]^ The changes in moisture content (thus breathing) could be detected by applying an 5V, 2kHz AC signal between the electrodes and measuring the resulting ionic current running through the textile which is proportionally related to the ionic conductivity of the substrate. Using portable wireless electronics (custom designed in our group, **Figure S11** and **S12**), we were able to capture respiratory activity from a volunteer and transmit the waveforms to a nearby mobile device running an Android app, developed also by our group (**Video S3**). With the embroidered respiration sensors, we could clearly detect different forms of respiratory activity including normal, paused, fast, shallow, slow and deep breathing without any post processing. By counting the peaks manually, the breathing rates (an important indicator of health and one of the four vital signs used in medicine^[33]^) for the different phases of respiratory activity could also be estimated.

#### Embroidered gas sensors for sensing NH_3_

For wearable sensing of gases, we produced embroidered gas sensors using cotton textiles as substrate (**Figure 4C**). Because NH_3_ has both diagnostic and environmental relevance, we tested the embroidered gas sensors for the detection of ammonia.^[36, 37]^ We produced two different types of chemiresistive gas sensors using PECOTEX with different underlying principles (or modes) of operation: *Sensor Mode I* consisted of a single continuous running stitch of PECOTEX and exploited the known phenomenon of dedoping of PEDOT when exposed to alkaline gases, which reduce the conductivity of PEDOT.^[38]^ *Sensor Mode II* had a similar geometry to the respiration sensor described above, consisting of two embroidered electrodes of PECOTEX, separated by the textile substrate. Using this geometry, we measure the ionic conductivity of the hygroscopic, cellulose-based textile substrate, the conductivity of which changes when exposed to water soluble gases such as ammonia (this sensing approach was reported by our group previously – see^[39]^ for more details).

Both Mode I Mode II sensors showed a response (that is change in conductivity *G)* when exposed to ammonia gas (Figure 4C): expectedly, while the conductivity of Mode I sensors decreased when subjected to 500 part-per-million (ppm) of ammonia, the conductivity of the Mode II sensors increased. The conductivity of Mode I sensors did not recover back to their initial baseline, hence showed a cumulative response as gaseous NH_3_ seemed to have modified the polymer permanently, whereas Mode II sensors exhibited a semi-reversible response - *i.e.,* when the flow of gas was shut-off, the conductivity of the sensor decreased but not completely back to the starting levels. We investigated the concentration dependent response of the Mode II sensors by exposing the sensors to varying levels of NH_3_ up to 1000 ppm (**Figure S13**). The sensors showed increasing response to higher concentrations in agreement with our previous work;^[37]^ however, when we repeated exposed the sensors to the same concentration of gas, we noted that the sensor response decreased (Figure 4C – inset). This is likely due to two competing processes of sensing of NH_3_ using Mode II sensors: while the conductivity of the PECOTEX embroidered electrodes decrease when exposed to ammonia in a non-reversible fashion, the ionic conductivity of the textile substrate increase reversibly resulting in a semi-reversible gas sensor the performance of which likely will degrade with use over time. Both Mode I and Mode II sensors are, therefore, likely most suitable for short-term use or will need to be regenerated by acid treatment after extended exposure to alkaline gases.^[40]^

## 3. Conclusion

In conclusion, we have produced an all-organic cotton-based conductive thread (PECOTEX) that is compatible with domestic and industrial computerized embroidery machines. PECOTEX reported in this work can be produced at sufficiently low-cost using roll-to-roll processing (**Table S1**): the cost of materials was 0.15 USD m^-1^ to produce low volumes of the conductive thread in our lab; we expect this number to decrease substantially when the production is scaled up industrially. PECOTEX exhibited superior characteristics compared to the commercially available Ag-based conductive threads including (**Table S2**): the average resistance after embroidery, number of breaks during embroidery, number of layers embroidered on top of each other *etc.* Furthermore, PECOTEX is compatible with computerized embroidery both in the top and bottom configuration as it can withstand the harsh mechanical conditions experienced during embroidery.

PECOTEX reported in this work have the following three disadvantages: *(i)* The electrical resistance of PECOTEX increase after embroidery up to 100 times. They remain, however, sufficiently conductive for most sensing applications.^[41–46]^ Furthermore, the electrical resistance of the embroidered patterns can be reduced by denser stitching, using a longer stich length or deposition of additional materials by electroplating.^[43, 47–49]^ *(ii)* In comparison to metal-coated conductive threads, the electrical resistance of PECOTEX were a few orders magnitude higher; however, the electrical resistance of the metallic threads increased and became comparable to the PECOTEX after embroidery. *(iii)* Although we did not study in detail, we did notice a drop in electrical conductivity of the patterns embroidered with PECOTEX over time. Interestingly the electrical characteristics of the threads stored without embroidery remained stable over time.

Although in this work, we have only shown the application of PECOTEX for wearable monitoring of cardiac, respiratory activity and gas sensing, the material presented is suitable for a large range of other applications. These include, but not limited to, wearable electrochemical sensors^[46, 50, 51]^, batteries^[52]^, heaters^[53]^, interconnects for wearable circuits and conductors for anti-electrostatic (ESD) clothing all of which can be produced at large volumes and cost-effectively through computerized embroidery.

## 4. Experimental Methods

### Pretreatment of cotton thread

Two-ply core-spun virgin cotton thread (JBS Olivia, South Korea) was desized by stirring in hot aqueous solution (80 ^o^C) containing 5 % wt. of Synthrapol (also known as METAPEX 38, Kemtex), which consists of isopropanol and ethoxylated and sulfated aliphatic alcohols, for 2 h to remove water soluble and oily sizes. The thread then was immersed in a 4:1 (vol:vol) aqueous bleaching solution for 5 h. The solution contained sodium hydroxide (scouring agent) and sodium hypochlorite (oxidative desizing agent) to oxidize residual organic contaminants removing the “cuticle” (waxy layer) from the cotton fiber. The threads were washed several times with DI water before finally oven-drying at 50 ^o^C for 1 h. The pretreatment efficiency was examined by applying water drops on the cotton thread. Successful pretreatment was confirmed with the formation of a hydrophilic thread.

### Roll-to-roll fabrication of the PECOTEX

We filled a 50 ml falcon tube with an aqueous solution of PEDOT:PSS (CLEVIOS PH 1000, produced by Heraeus), ethyleneglycol (EG) and divinylsulfone (DVS) with varying ratios. EG and DVS were purchased from Sigma-Aldrich and used without any further modification. The cotton thread was first wound on a bobbin manually and pulled through four falcon tubes all containing the same PEDOT:PSS mixture using a servo-motor (Parallax Standard Servo) before winding on another bobbin after coating. The speed rotation of the servo-motor was controlled by an Arduino Nano microcontroller board which was programmed using the Arduino IDE, installed on a Windows PC, over a USB connection. The optimum speed was found to be 3.6 m min^-1^. After the thread was coated, before winding on the bobbin, it was briefly dried by a hairdrier to remove excess solvent to prevent sticking.

### Computerized embroidery

Embroidery designs were made on a Windows PC using the PREMIER+™ 2 software. After the designs were completed, they were downloaded onto the computerized embroidery system (DESIGNER DIAMOND Royale™ by HUSQVARNA VIKING) before stitching. After the fabric substrate was placed in the hoop and inserted into the embroidery machine, the software design was embroidered on the substrate, lasting typically a few minutes.

### Raman spectroscopy

Resonant Raman spectroscopy was performed on the sample using a Renishaw inVia microscope in a back-scattering configuration at an excitation wavelength of 633 nm with 10% of 12 mW laser power under 50× magnification. Samples were placed directly under the laser and focused to generate spectra. Multiple datasets of each sample were used to obtain an average representative spectrum for all normalized Raman spectra.

### Microstructural characterization

Scanning electron micrographs were taken using JEOL, JSM-6010LA at 20 kV electron beam energy. The optical micrographs were obtained by a Brunel SP202XM metallurgical microscope. A Nikon D3200 camera was attached to the microscope for the acquisition of the images using DigiCamControl open-source software. For focus stacking to improve the limitation of depth-of-field, multiple images were taken at different focus and stacked together using Helicon Focus software.

### Mechanical and electrical testing

The mechanical and electromechanical tests were carried out using a MultiTest 5-xt tensile testing machine controlled by Emperor Force software (Mecmesin) and 72-7730A source multimeter from TENMA. For mechanical testing, the thread samples were strained at a speed of 50 mm min^-1^ until complete fracture. The (electro)mechanical tests were performed using PECOTEX samples of 10 cm in length. For electromechanical testing, PECOTEX samples were connected to the multimeter using crocodile clamps on both ends to measure their electrical resistance during the test. The PECOTEX samples were subjected to a cyclic force between 1 and 2.5 N. The electrical resistance of all samples was measured using the same multimeter as described above (*i.e.* 72-7730A from TENMA). The IV curves of PECOTEX were acquired using a Keithley 2450, Source-meter (Figure S8).

### Electrochemical testing

Cyclic voltammograms were acquired using a handheld potentiostat (PalmSens3, PalmSens BV, The Netherlands) with the supplied PSTrace 5.3 software in a three-electrode electrochemical cell configuration. A commercial screen-printed drop sensor (supplied from Metrohm DropSens, Switzerland) was used to conduct the measurements with 3 cm of PECOTEX attached to the strip as the working electrode, with printed Ag/AgCl as reference and C counter electrode. The scanning windows for the applied potential was set to −1.0 - 1.2V with varying sweep rates. All electrochemical tests were performed in an aqueous solution of 0.1 m Potassium chloride (KCl from Sigma Aldrich) and 20 mM ferrocyanide (K_3_[Fe(CN)_6_] from Sigma Aldrich).

### Washability experiments

The washability tests were conducted following the American Association of Textile Chemists & Colorists protocol by using a domestic laundry machine (Candy GOW475 Washer Dryer): The cotton fabrics embroidered with PECOTEX were placed into a washing bag and placed in the washing machine along with 1.8 kg of fabric serving as a ballast. The washing cycle lasted 60 mins in total: 50 mins of washing in water on hand-washing mode and 10 mins of mechanical drying at 600 RPM and 30 ^o^C. The electrical resistance measurements were taken after drying the samples for 1 h at 50 ^o^C in an oven in air. In order to confirm sufficient drying, one of the samples was left overnight before repeating measurements the next day which matched the initial readings (after 1h at 50 ^o^C).

### ECG experiments

The ECG recordings were obtained from a volunteer using an ESP32-based microcontroller board attached to an AD8232 SparkFun Single Lead Heart Rate Monitor. The waveforms captured were transferred to a PC running MATLAB over WiFi. The waveforms collected were also post-processed in MATLAB (see supporting information Figure S9 and S10 for more information on post processing). The disposable commercial ECG pads were sourced from Amazon. The ECG cable was purchased from SparkFun. Conductive gel (Elektroden-Gel) was purchased from Amazon.

### Respiration sensing experiment

For data acquisition, a custom-made read-out board based on ESP32 was used. An alternating voltage was applied across the sensing electrodes and an opamp (AD820)-based transimpedance amplifier was used to estimate the impedance of the sensors. The range of amplification was adjusted by switching between different gain resistors using a multiplexer (MAX335). The waveforms captured by the read-out board was transmitted to a nearby android phone in real-time via Bluetooth Low Energy (BLE) which contained a homemade app. See **Figure S11** and **S12**.

### Ammonia sensing experiment

A homemade gas sensor characterization setup was used to adjust the concentration of ammonia in the sensing environment using three mass flow controllers (MFCs, type GM50A from MKS) while measuring the electrical conductance using a custom-board. For more information on the gas sensor characterization setup used see our previous publication.^[36]^

All the human experiments were risk assessed.

## 5. Supporting Information

Supporting Information is available from the Wiley Online Library or from the author.

## 6. Acknowledgments

We would like to thank EPSRC (EP/G037515/1, EP/L016702/1, Imperial Impact Acceleration Account), Cytiva, Imperial College London, Department of Bioengineering, Bill and Melinda Gates Foundation (Grand Challenges Explorations scheme under grant number: OPP1212574) and the US Army (U.S. Army Foreign Technology (and Science) Assessment Support program under grant number: W911QY-20-R-0022), Wellcome Trust (Grant No. 207687/Z/17/Z) and Innovate UK (10004425). H-S.L. thanks ESRC LISS DTP (2453729) and F.A. thanks King Abdulaziz City for Science and Technology and the Saudi Ministry of Education. F.A is also grateful for Helle and Karen and their lovely shop ‘The Sewing Rooms’ for providing support and guidance in all computerized embroidery and textiles related matters.

Received: ((will be filled in by the editorial staff))

Revised: ((will be filled in by the editorial staff))

Published online: ((will be filled in by the editorial staff))

