## Supporting Information for "PEDOT:PSS-Modified Cotton Conductive Thread for Mass Manufacturing of Textile-Based Electrical Wearable Sensors by Computerized Embroidery"

† Significant contributor

* Corresponding Authors

F. Alshabouna,

Dr. F. Güder,

**Keywords:** Industry 4.0, Conductive Cotton Thread, Computerized Embroidery, Wearable Sensors, Electrocardiography, Respiration Monitoring, Gas Sensing

**Video S1** Slow Motion Video of Embroidery of Interdigitated Electrodes for Respiration Sensor.

**Video S2** Real-Time Electrocardiography with Embroidered Sensors on a T-shirt (Commercial Electrodes vs Embroidered Electrodes).

**Video S3** Respiration Monitoring with Embroidered Sensors in a Disposable Mask.

**Table S1**. Materials cost used to produce 100 m of PECOTEX

| Material | Quantity | Cost (USD) | Quantity to produce 100m of thread (USD) | Cost to produce 100m of thread (USD) |
|---|---|---|---|---|
| **Bleach** | 1000 ml | 0.86 | 300 ml | 1.5 |
| **Metapex** | 250 ml | 5.25 | 5 ml | 0.11 |
| **PEDOT:PSS** | 1000 ml | 452 | 20 ml | 9.04 |
| **DVS** | 100 g | 416.29 | 1 g | 4.16 |
| **EG** | 2000 l | 160.93 | 0.6 g | 0.05 |
| **Cotton thread** | 4000 m | 5.35 | 100 m | 0.13 |
| **Total** | | | | 15.0 USD/100m |

**Table S2**. Embroidery performance of PECOTEX and commercial conducive thread Shieldex® of an O-shape embroidered pattern (top thread configuration). Shieldex® plated-silver yarns were extremely fragile and broke during handling and winding onto bobbin.

| Criteria | PECOTEX | Shieldex |
|---|---|---|
| **Material** | All organic (PEDOT: PSS-modified cotton) | Silver-plated polyester |
| **Average resistance after embroidery** | 0.621 k Ω/cm | 0.758 k Ω/cm |
| **Number of snaps during a single machine embroidery** | - | 3 |
| **Thread winding onto bobbin** | Automatic using bobbin winder on embroidery machine. | Hand-winding |
| **Multi-Layers feasibility** | Electrical resistance decreased with number of layers (up to 3 layers). | After second layer, electrical resistance increased beyond multimeter limit. |

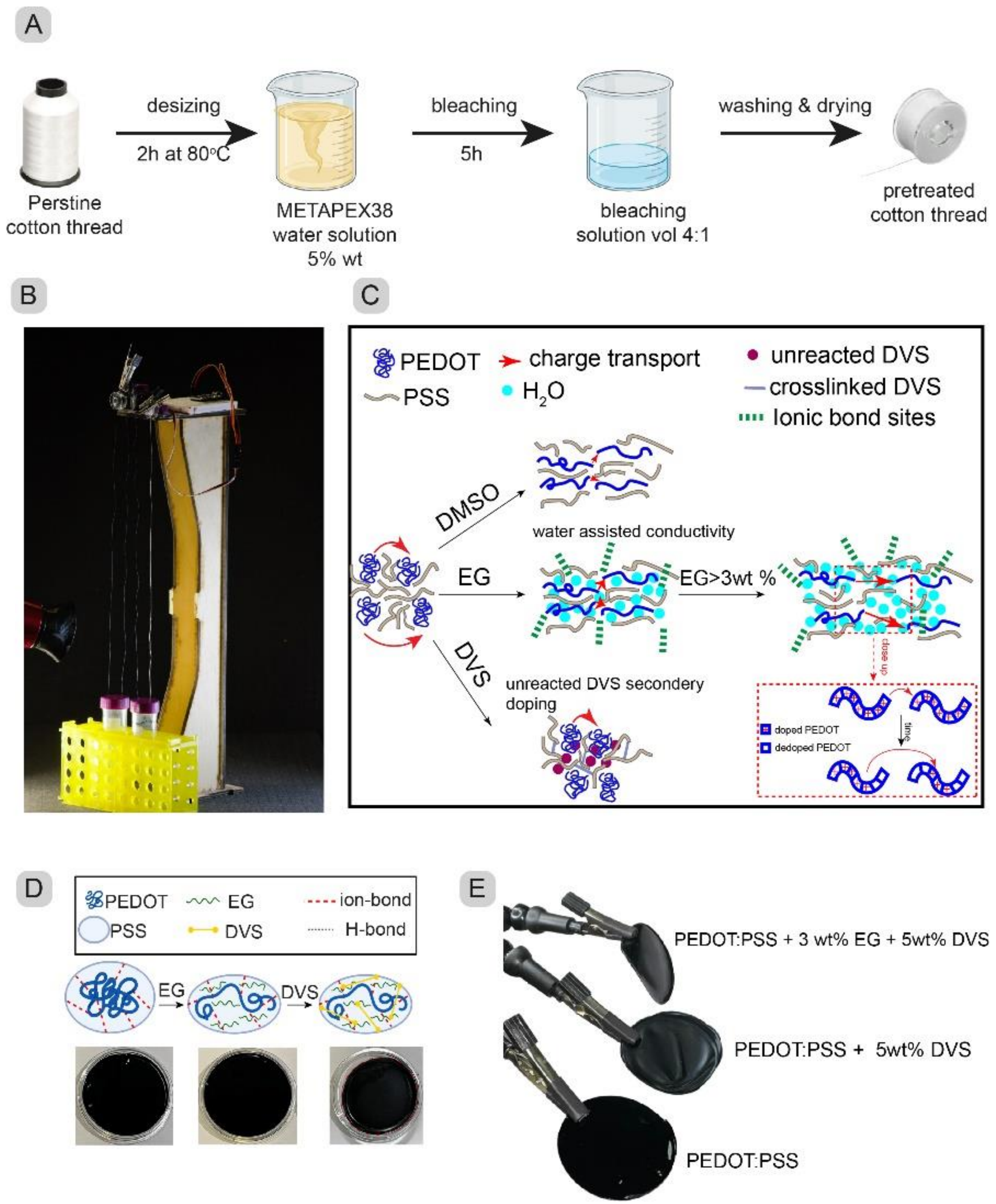


**Figure S1. (A)** Process flow for pretreatment of virgin cotton threads. **(B)** Picture of roll-to-roll setup used in manufacturing PECOTEX. **(C)** Schematic diagram of the effect of additives on the molecular conformation of PEDOT:PSS and subsequent effect on charge transport mechanism.

**(D)** Pictures of PEDOT:PSS freestanding films produced in Petri dish: alone, with EG 3wt%, and with both EG 3wt% and DVS 5% (left to right) **(E)** Pictures of freestanding films removed from Petri dish showing the effect of each additive on the film flexibility. The addition of DVS

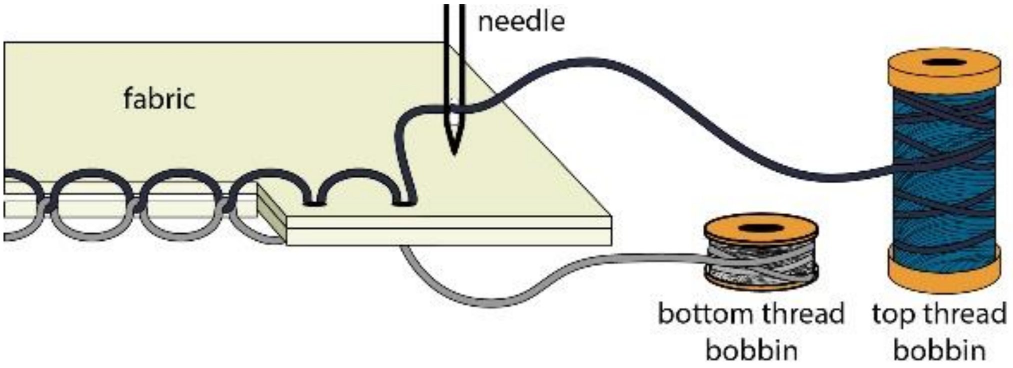


**Figure S2.** Schematic illustration of embroidery process. The top thread is accommodated in the eye of the needle, the needle penetrates the fabric through to the opposite side, where a revolving hook rotates the top thread around the bottom thread, forming a knot (lock stitch).

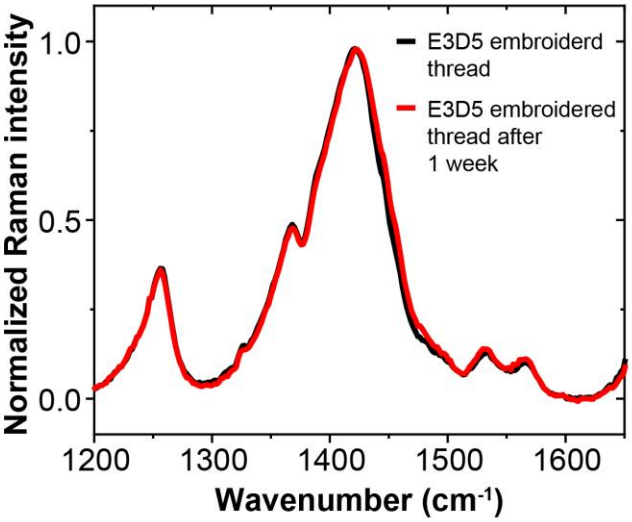


**Figure S3.** Normalized Raman spectra of embroidered conductive thread (E3D5 formulation) recorded after stitching and after 1 week.

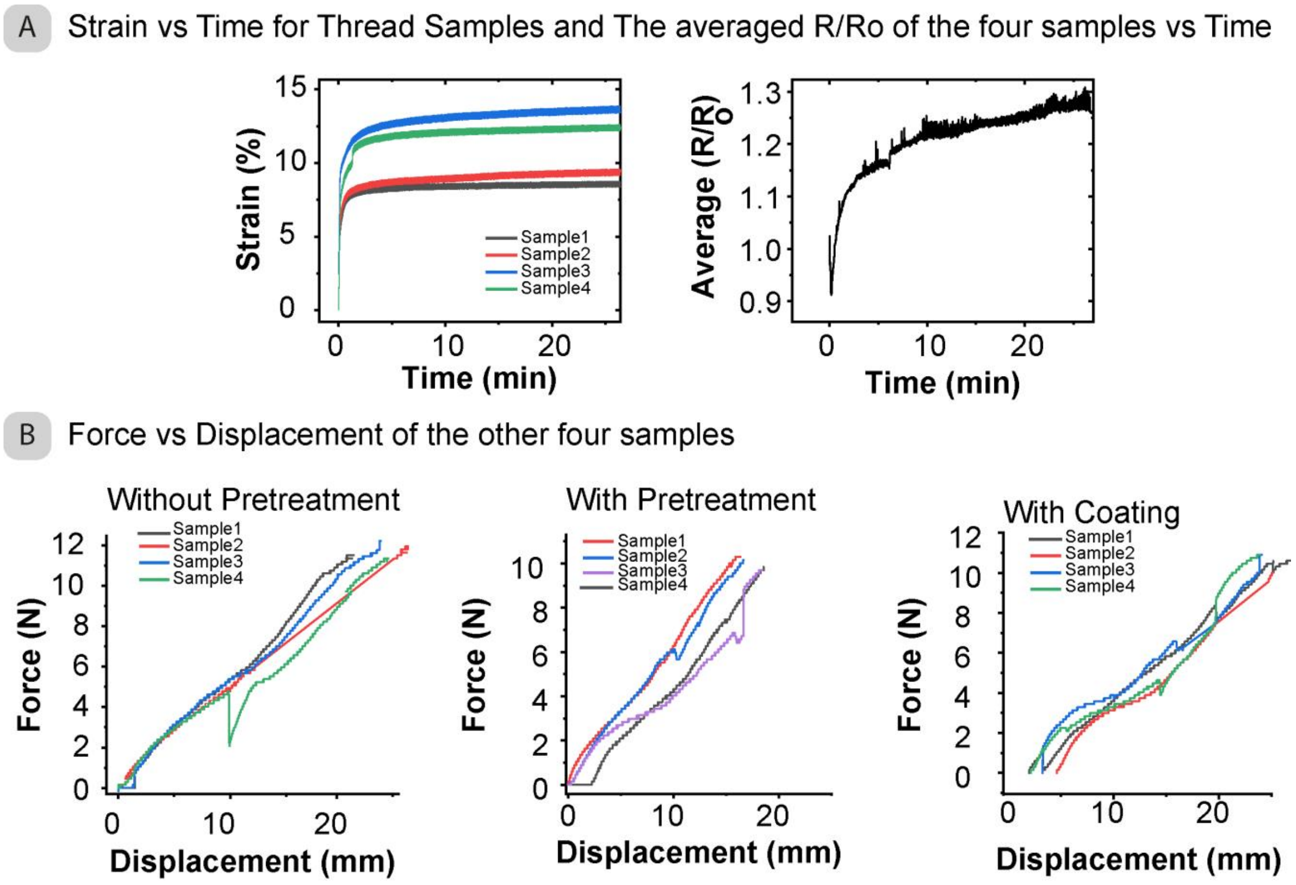


**Figure S4. (A)** Change in resistance (averaged) under cyclic strain for the rest of the samples (n=4). This shows that the maximum resistance change for all the samples was less than 40%. **(B)** The tensile test shows force vs strain of the rest of the samples thread ( n=4 each).

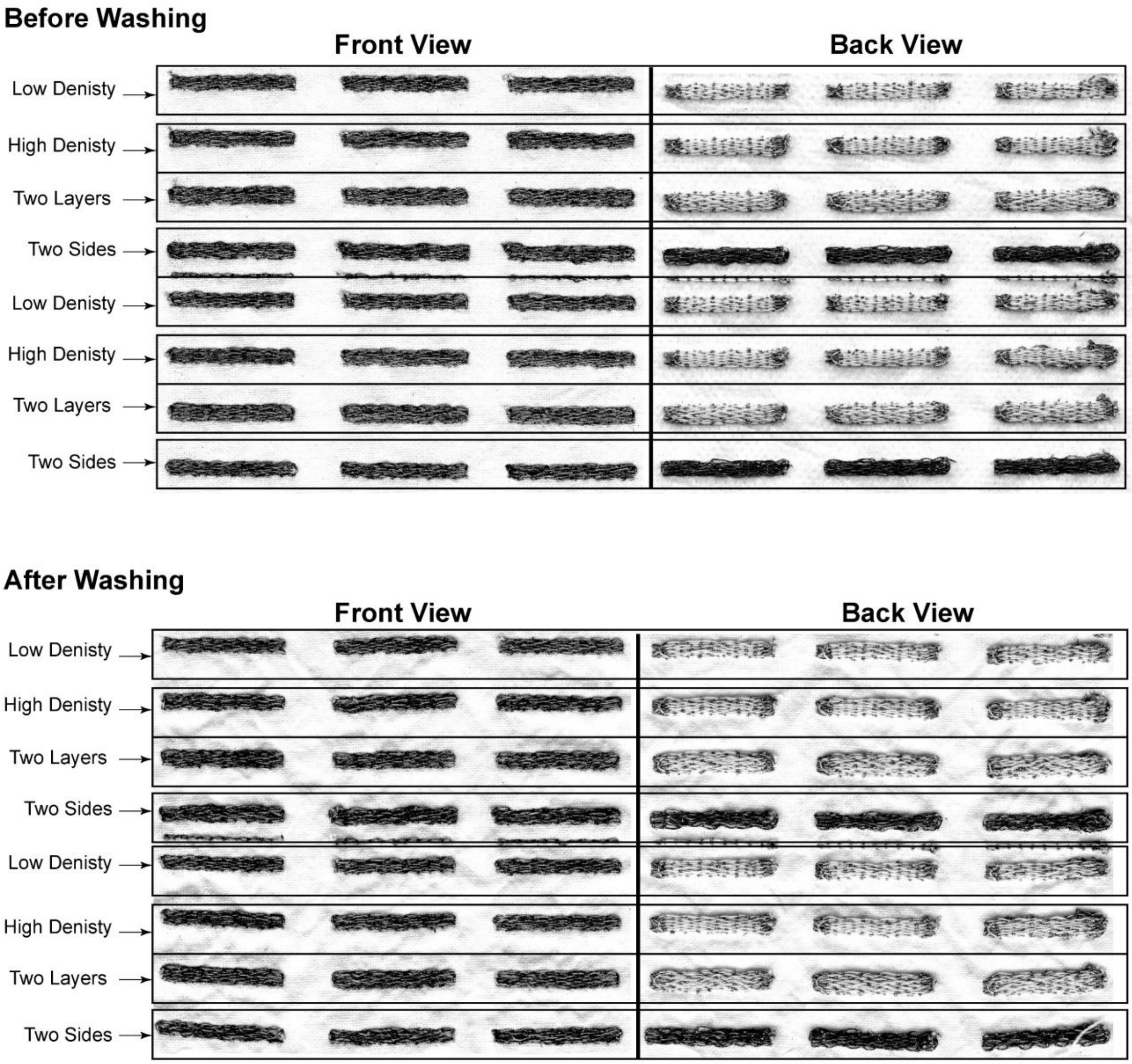


**Figure S5**. Scanned pictures of front and backside of embroidered patterns of PECOTEX (20 × 2 $mm^2$) with different stitch densities, multilayers of stitching and combining top and bottom bobbin threads configurations, before washing cycles (top) and after washing cycles (bottom).

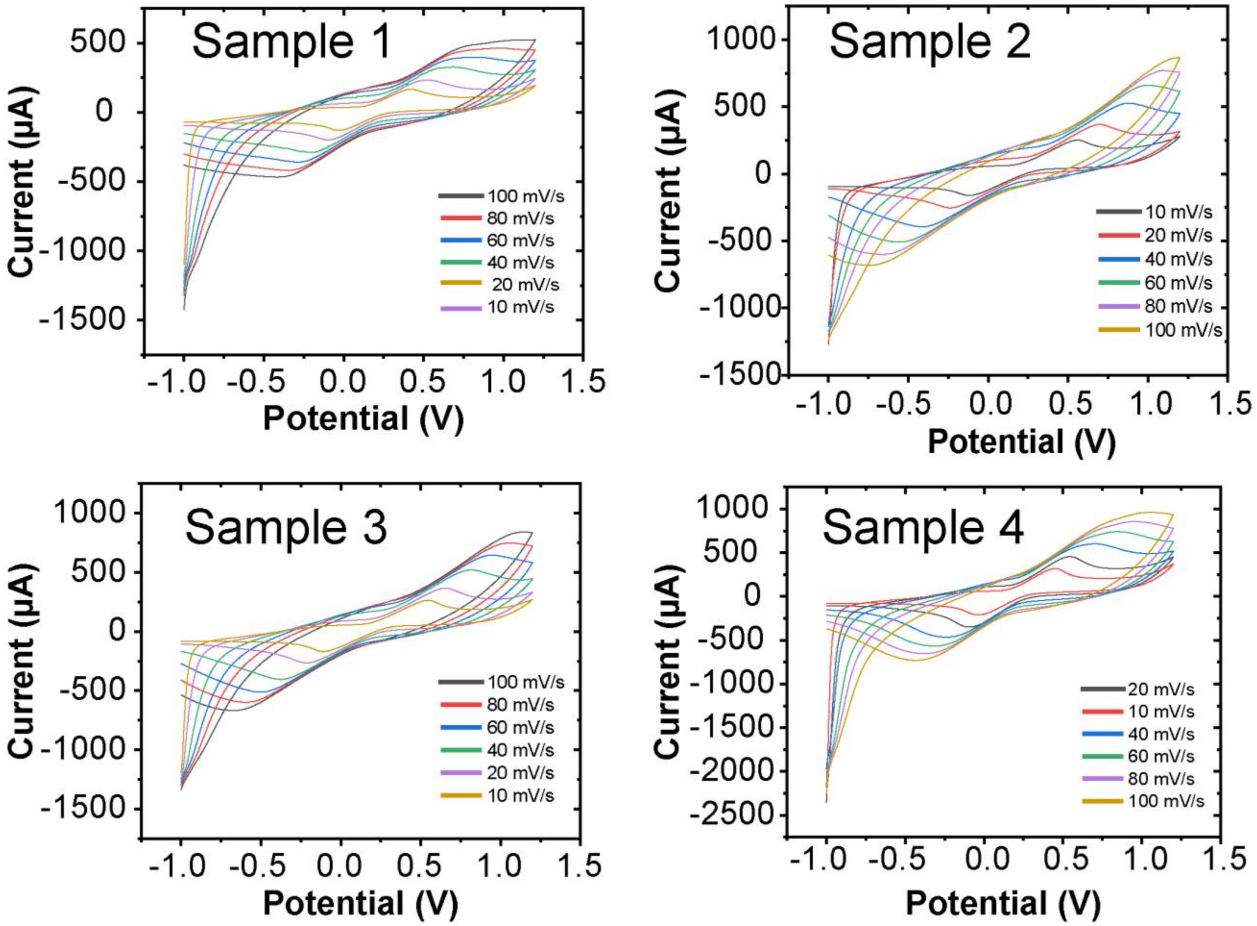


**Figure S6**. The cyclic voltammogram of the rest of PECOTEX samples (3 cm long) (total n = 4) at the scan rate from 10 m V $s^{-1}$ – 100 m V $s^{-1}$ in 20 m M Ferrocyanide/0.1 M KCl solution.

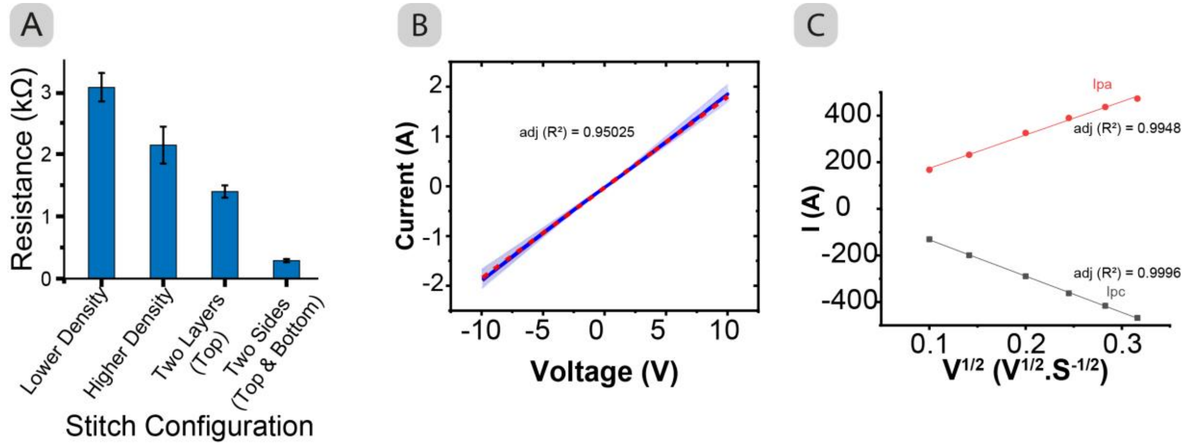


**Figure S7**. **(A)** Variation in electrical resistance of embroidered patterns of PECOTEX with different stitch densities, multilayers of stitching and combining top and bottom bobbin threads configuration. **(B)** IV curve for PECOTEX (n=4, 4 runs per sample), showing typical liner resistor behavior. **(C)** Relationship between the peak current ($i_p$) and the square root of scan rate ($v^{1/2}$) in cyclic voltammetry of PECOTEX.

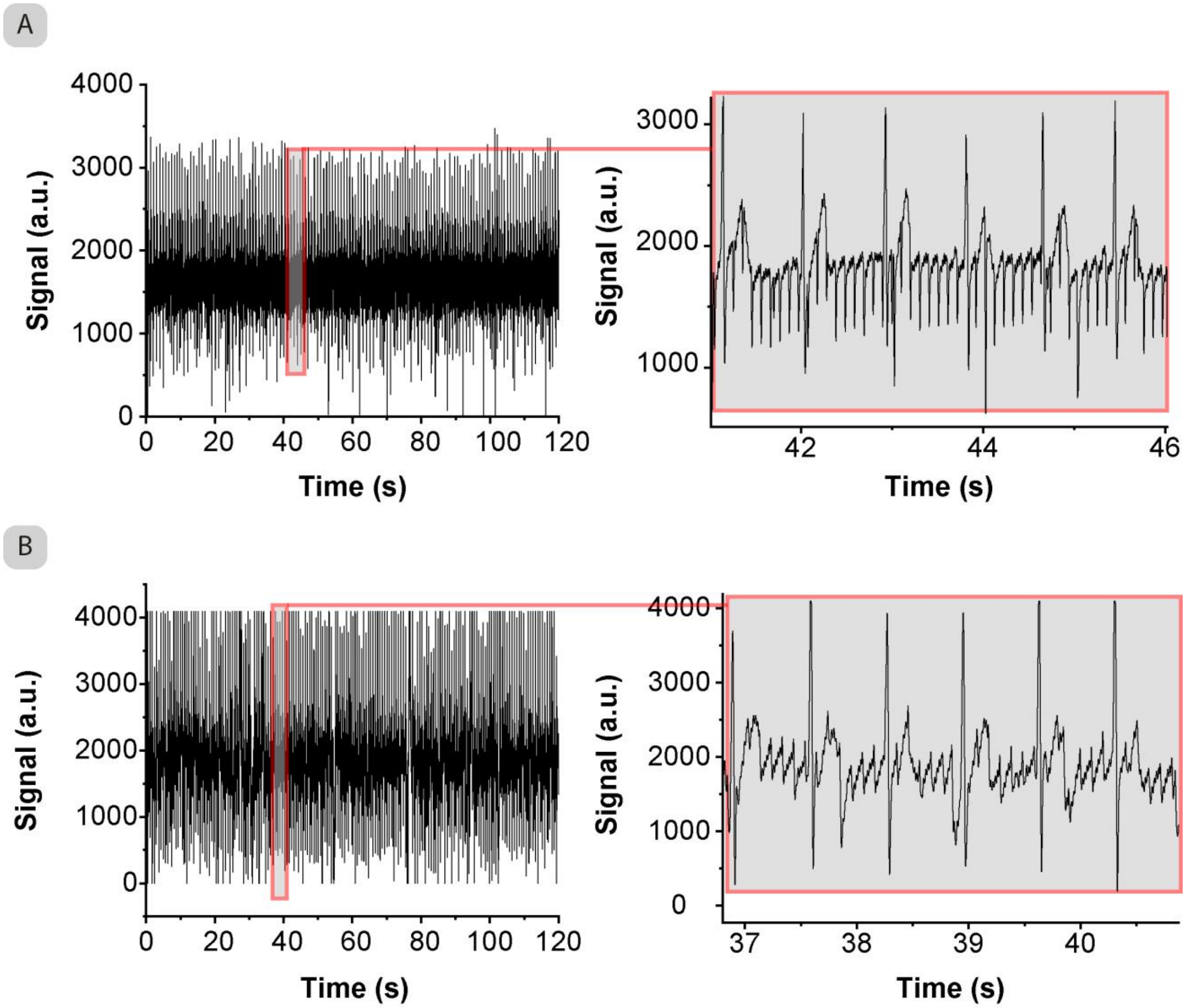


**Figure S8**. **(A)** Raw ECG recordings of cardiac signals of PECOTEX stitched electrodes on a t-shirt for two minutes (left) and a selected 6 cycles (right). **(B)** Raw recordings of cardiac signals of commercial electrodes for two minutes (left) and a selected 6 cycles (right).

**Post-processing of ECG signals**: We powered the microcontroller using a USB cable through the computer for the recording of ECG signals at a sampling rate of 100Hz. The AC power-line contamination effected the quality of the readings because of the alternating current oscillations at 60Hz. A notch filter is commonly used for the elimination of power-line noise. Similarly, after taking the signal to the frequency domain using “fft” function of MATLAB, we observed a periodic noise at 1Hz as seen in Figure S8. To eliminate this periodic noise, we used “iircomb” function of MATLAB, a specific version of an Infinite impulse response (IIR) filter. In the filter design, we set the following parameters: sampling rate (Fs) as 100Hz, unwanted periodic frequency (Fo) as 0.97Hz and a quality factor (Q) as 15. “iircomb” function requires filter order (Fs/Fo) as an integer. Therefore, we needed to round (100/0.97) to 103 and adjust the quality factor to make sure the bandwidth of the filter covers the unwanted notches. Finally, we implemented a Savitzky–Golay filter to smooth the signal without distorting the ECG waves using “smooth” function of MATLAB.

Here is the implemented function.

```
Fs = 100; Fo = 1; Q = 15; BW = (Fo/(Fs/2))/Q;
[b,a] = iircomb(round(Fs/Fo),BW,'notch');
fdata = filtfilt(b,a,data);
fdata2 = smooth(fdata,30,'sgolay');
```

As seen in **Figure S9**, unwanted peaks in the raw ECG data are eliminated after the implementation of notch and smoothing filters and ECG waves can easily be recognized.

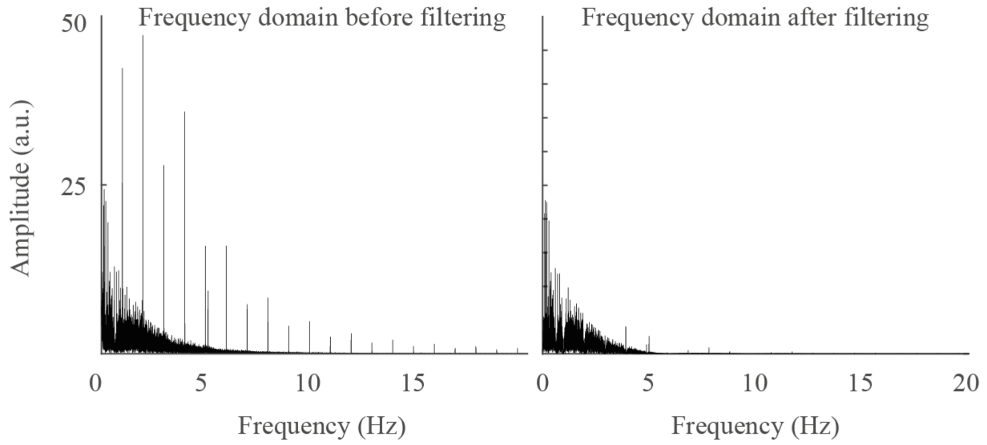


**Figure S9**. Frequency domain of the signal before and after implementing the filters. The periodic noise in the raw data is eliminated by implementing a notch filter with a frequency of 1Hz.

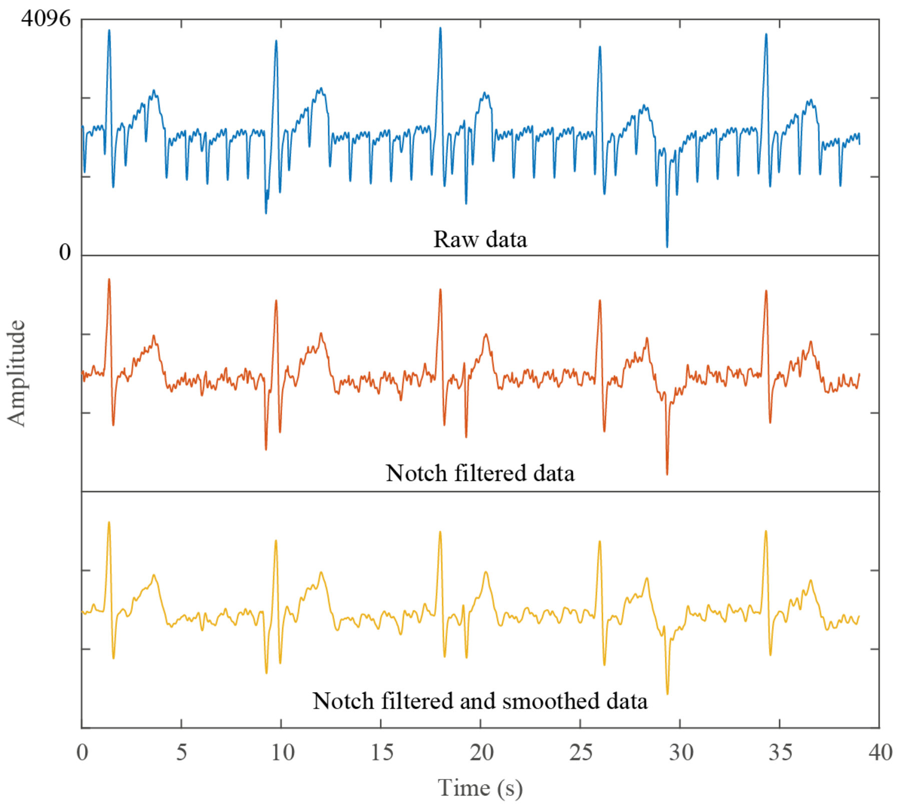


**Figure S10**. Removing noise from the raw data by implementing a notch and smoothing filter successively.

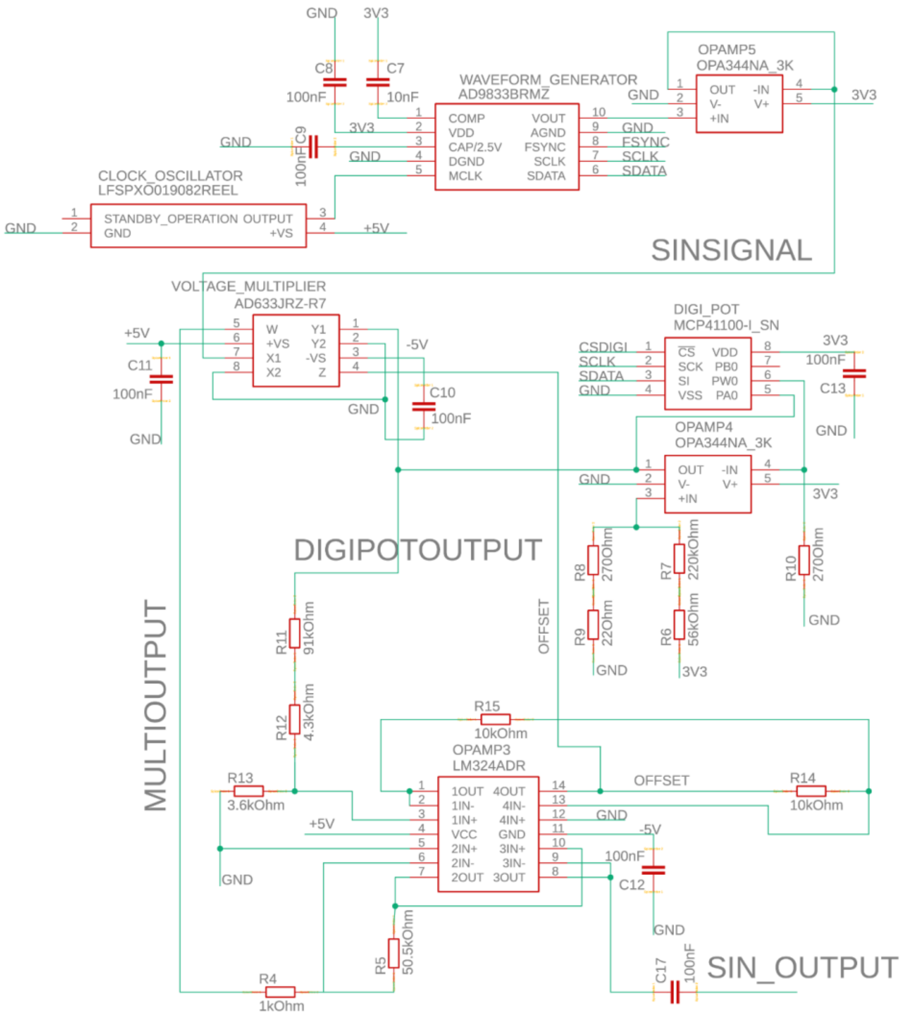

**Figure S11**. This circuit creates the sinusoidal wave to measure impedance in the textile respiration sensor. A 24MHz clock oscillator clocks a waveform generator (AD9833) which is fed into a voltage follower (OPAMP5) to create the base signal ‘SINSIGNAL’ in a frequency between 10Hz-10kHz. The DC offset of the sinusoidal signal is then removed using a voltage multiplier (AD633) and amplified in a transimpedance amplifier configuration using an adjustable gain (digital potentiometer, MCP41100) to reach the desired voltage range (max. ca. +/- 4.2V). The final signal ‘SIN_OUTPUT’ is the input to the textile respiration sensor.

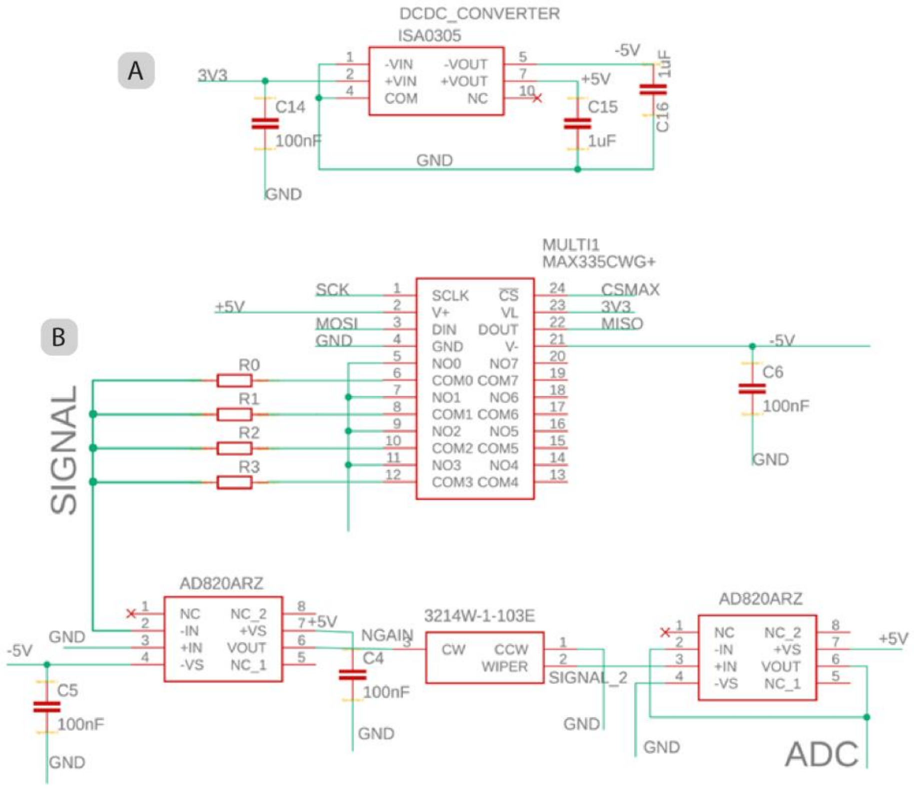


**Figure S12**. **(A)** A DC-DC converter is used to extend the 3.3V from a standard Arduino to -5 / +5 V to achieve proper AC sinusoidal waveforms (see Figure 10). **(B)** This circuit takes the sensor signal as an input, amplifies, and feeds the signal into the Arduino ADC. The sensor input is ‘SIN_OUTPUT from Figure 10 and the output from the sensor (‘SIGNAL’) is amplified using a transimpedance amplifier configuration. A multiplexer (MAX335) chooses the correct gain resistor (50M, 10M, 1M, 100k) and a potentiometer (3214W) is used to downscale the 10Vpp signal range to the 3.3V to be read by the Arduino ADC.

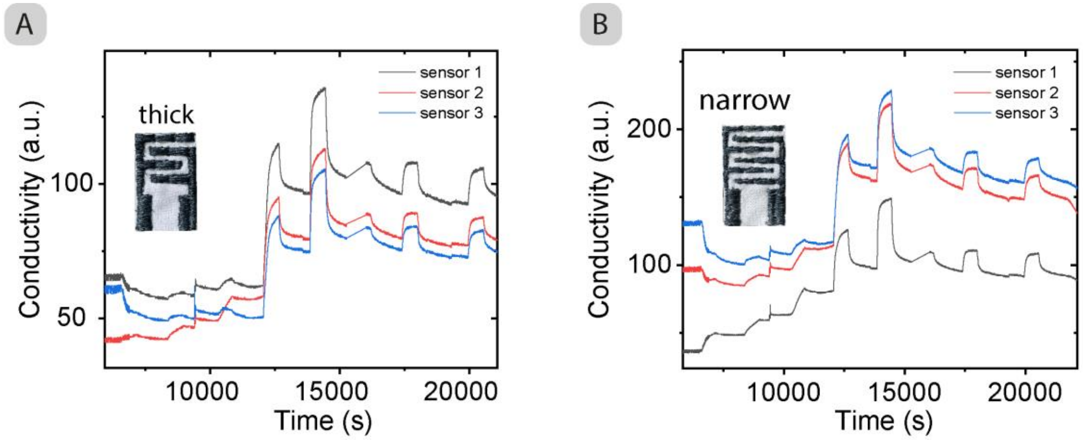


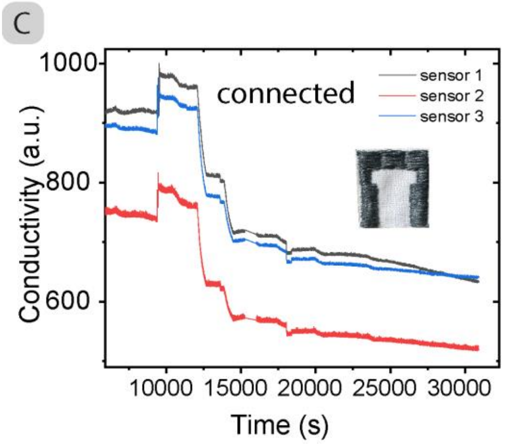


D

| $NH_3$ Concentration (ppm) | Time in (s) | Time out (s) |
|---|---|---|
| 5 | 6629 | 7229 |
| 10 | 8429 | 9629 |
| 50 | 10229 | 12029 |
| 500 | 12029 | 14429 |
| 1000 | 13829 | 16829 |
| 100 | 15629 | 19229 |
| 200 | 17429 | 21629 |
| 200 | 19229 | 24029 |

**Figure S13**. **(A-C)** Change in conductivity response of three types of textile-based sensor during $NH_3$ exposure, digitally smoothed to 100 points. **(D)** Summary table of $NH_3$ concentration and on/off times.
